# Long-term CRISPR array dynamics and stable host-virus co-existence in subsurface fractured shales

**DOI:** 10.1101/2023.02.03.526977

**Authors:** Kaela K. Amundson, Simon Roux, Jenna L. Shelton, Michael J. Wilkins

**Affiliations:** Colorado State University, Fort Collins, CO, USA; DOE Joint Genome Institute, Lawrence Berkeley National Laboratory, Berkeley, CA, USA; United States Geological Survey, Reston, VA, USA

## Abstract

Viruses are the most ubiquitous biological entities on earth. Even so, elucidating the impact of viruses on microbial communities and associated ecosystem processes often requires identification of strong host-virus linkages – an undeniable challenge in many ecosystems. Subsurface fractured shales present a unique opportunity to first make these strong linkages and subsequently reveal complex long-term host-virus dynamics and trends in CRISPR array size and frequency. Here, we sampled two replicated sets of fractured shale wells for nearly 800 days (Denver-Julesburg Basin, Colorado, USA). We identified a relatively diverse microbial community with widely encoded CRISPR viral defense systems, which facilitated 2,110 CRISPR-based viral linkages across 90 host MAGs representing 25 phyla. Leveraging these linkages with timeseries data across differing well ages, we observed how patterns of host-virus co-existence develop and converge in this closed ecosystem through time. We observed a transition to smaller CRISPR arrays in older, more established wells, potentially reflecting a natural progression where CRISPR arrays harbor fewer, yet more effective spacers that target viral genes with fewer mutations. Together, our findings shed light on the complexities of host-virus temporal dynamics as well as possible drivers of spacer loss and acquisition within CRISPR arrays of diverse microbial populations through time.

## Introduction

Viruses are abundant and important constituents of microbial communities in nearly all ecosystems. Consequently, bacteria and archaea, like all living things, are subject to near constant threat of viral predation. In response, many bacteria (~40-60%) and archaea (~90%) deploy CRISPR-Cas viral defense systems (1–4). CRISPR works by recording memories of viral interactions via integration of small pieces of viral DNA (‘spacers’) within the hosts’ CRISPR array, that are interspaced with identical repeat sequences and flanked by Cas (CRISPR-associated) genes (5–13). These saved memories help to protect the host against recurrent invasion by the same viral population by more rapidly identifying and degrading the invading nucleic acids, analogous to antibodies in the human immune system (5–7,10,12,14).

Spacers within CRISPR arrays therefore provide a record of past interactions between a host and viral population, and host-viral linkages can be made by matching the hosts’ CRISPR spacers to protospacers in viral genomes (11,15–27). However, the presence of CRISPR systems within the microbial community is often a limiting step to making strong host-virus connections; CRISPR defense is most likely advantageous in ecosystems where host and viral populations repeatedly interact, such as environments dominated by biofilms or those hosting lower microbial and viral diversity (28–30). Additionally, CRISPR has been shown to be more widespread in some ecosystems relative to others, such as anoxic environments or those with elevated temperatures (14,28,31–33).

Despite the important role of CRISPR in viral defense, little is known about the drivers of array size and frequency, and how these might change through time for host populations within natural microbial communities. Successful incorporation of a spacer should provide the host future defense upon interaction with the same viral population. However, CRISPR arrays do not grow exponentially as spacers can indeed be lost, (34–37), and host-viral co-existence despite CRISPR has been observed (38). Additionally, spacers nearest to the leading end of the CRISPR arrays are most likely to be effective, as they typically represent more recent viral interactions with less time for mutations to occur within the viral protospacer, although recombination can also influence CRISPR array architecture (8). Thus, it has been hypothesized and shown in laboratory experiments that select spacers may be more favorably retained if they target evolutionarily conserved portions of the viral genome, providing more effective long-term viral defense (13,39,39). Although ecosystem resources and genome size are not necessarily limiting factors to array size (40,41), other studies have modeled the optimum CRISPR cassette size based on other factors, such as viral diversity and tradeoffs between Cas machinery and array size (42–44). As a result, it has been suggested that maintaining smaller arrays, on the order of a few dozen to a hundred spacers, may be the optimal size for CRISPR arrays that provide broad protection against a range of viruses but do not overwhelm CRISPR machinery (42,44,45). However, many of these insights are derived from modeling or laboratory experiments and there remains a need to observe similar patterns in natural ecosystems.

To address this knowledge gap, we used a temporally-resolved dataset from six subsurface fractured shale wells to interrogate host-virus dynamics and CRISPR arrays in a natural ecosystem. Subsurface fractured shales, which are relatively closed ecosystems with limited immigration, elevated temperatures, lower microbial diversity and likely dominated by biofilms, present an opportunity to address these questions through strong CRISPR-based host-viral linkages (21,23,46–49). We hypothesized and found that CRISPR viral defense systems were widely encoded across hosts within shale microbial communities. Building on this, we applied multiple bioinformatic approaches to identify CRISPR spacers in both recovered host genomes and metagenomes and made strong host-virus linkages for many of the recovered host genomes. This approach also facilitated investigations into CRISPR array size and frequency and revealed contrasting patterns between hosts in older, established shale wells relative to younger, newly established wells. Our results provide additional evidence supporting the hypothesis that smaller CRISPR arrays may be more effective for optimal host defense against viral predation, while larger CRISPR arrays may be the temporary result of an ecosystem perturbation leading to new virus-host encounters. To our knowledge, our study represents one of the most extensive analyses of long-term, host-viral temporal dynamics with CRISPR-based linkages in a natural ecosystem to date.

## Results & Discussion

### Fractured shale ecosystems provide a unique opportunity to investigate virus-host temporal dynamics

We sampled fluids from two distinct sets of hydraulically fractured oil & gas wells in the Denver-Julesburg (DJ) Basin for nearly 800 days (Colorado, USA). The two sets of wells were defined by their age relative to the initial fracturing process: the ‘established’ wells operated for nearly three years prior to the initiation of our sampling campaign (DJB-1, DJB-2, DJB-3), while we began sampling the ‘new’ wells shortly after they had been hydraulically fractured (DJB-4, DJB-5, DJB-6) (Figure S1). All three wells within each group were located on the same frack pad and subject to the same drilling and hydraulic fracturing process, resulting in three replicate wells for each group.

From a total of 78 metagenomes across all six wells (Figure S1 & Table S1), we recovered 202 unique metagenome assembled genomes (MAGs) representing 29 phyla and 2,176 unique viral MAGs (vMAGs) from the subsurface communities. Many of the dominant and persisting MAGs – encompassing bacterial taxa affiliated with *Clostridia, Thermotogae, Fusobacteriia*, and *Synergistia* and archaeal taxa affiliated with *Methanosarcinia, Methanomicrobia*, and *Thermococci* (Figure S2) – have been reported in other engineered subsurface environments (21,50–54), and their relative abundances in this system reflected patterns observed in complementary 16S rRNA gene analyses (Figure S3).

Microbial communities, however, are not static through time. The spatial structure and taxonomic composition of the persisting microbial communities in these subsurface fracture networks likely develops as the ecosystem ages. Taxa unable to tolerate high temperatures and elevated salinity are likely outcompeted, while biofilms and spatially distinct niches likely emerge and expand (55,56). Thus, we expect that microbial communities within the established wells are more spatially heterogeneous and partitioned into more stabilized niches, while microbial communities in the new wells are initially well mixed, more spatially homogenous, and lack established biofilms (49,57).

In agreement with these assumptions, we observed higher host (bacterial and archaeal) and viral alpha diversity in the established wells relative to the new wells (*Wilcox, p=5.563e-06*) (Figure S4). Alpha diversity also generally increased through time in all wells, likely reflecting the development of niches fostering more diverse taxa (Figure S5). Notably, microbial communities from the new wells became more similar to those in the established wells over time (Figure S6). Although community composition did indeed fluctuate, host and viral community dynamics generally mirrored one another – suggesting viral predation is occurring (SI & Figure S7). Indeed, host-viral dynamics in the older, established wells were generally more strongly correlated than viral-host dynamics in the new wells (*Spearman Rho: established wells =0.87, 0.87, 0.44; new=0.51, 0.71, 0.77*) (Figure S7). Together, these results illustrate the temporal juxtaposition of the two sets of wells, yet the connectedness of host-viral dynamics in increasingly diverse microbial communities within a closed, subsurface ecosystem.

### Genome-resolved analyses reveal abundant evidence for widespread CRISPR viral defense systems in fractured shales

CRISPR viral defense systems were widely encoded across the DJ Basin microbial communities. In total, 123 of our 202 MAGs (~60%) spanning 25 of the 29 phyla contained a detectable CRISPR array (Figure 1 & Table S2). We identified CRISPR arrays in a higher proportion of MAGs from the established wells (67%) relative to the new wells (54%), potentially indicating selection for microorganisms that encode CRISPR as a viral defense system in an environment where viral predation is likely recurrent. The high proportion of MAGs that contained a CRISPR array is not unexpected, as CRISPR systems are more widely encoded in closed ecosystems, biofilms, and ecosystems with elevated temperatures – three characteristics of fractured subsurface shales.

**Figure 1.**
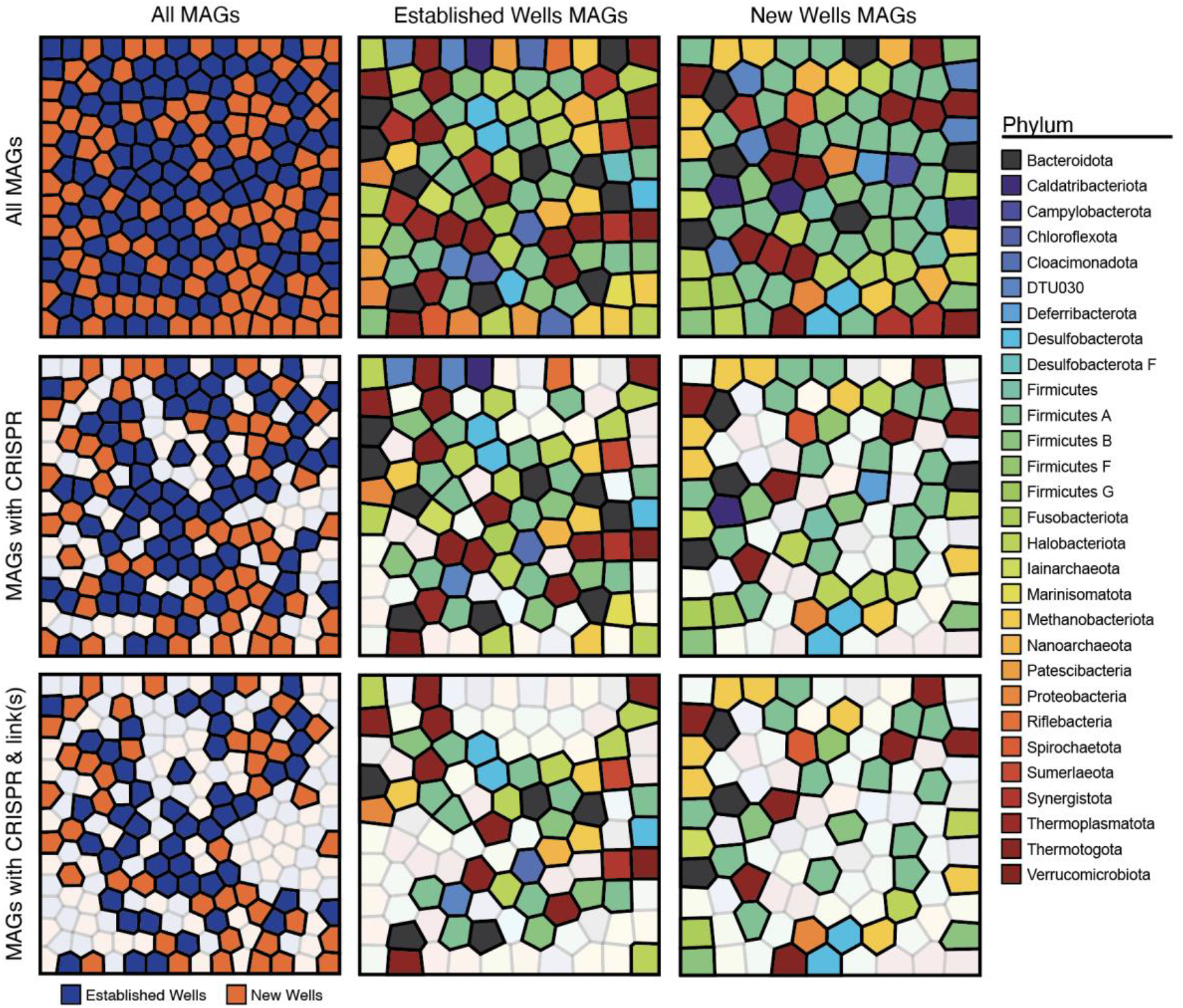
Host (bacterial & archaeal) MAGs recovered from this dataset. Each polygon represents an individual MAG. Columns illustrate the database of recovered host genomes: first by all MAGs colored by the well group from which the MAG was recovered (left column), then MAGs are colored by phylum for the established wells (middle column) and new wells (right column). Rows illustrate which MAGs had a detectable CRISPR array (middle row) and CRISPR and viral linkage(s) (bottom row) with lighter colored polygons representing MAGs that did not encode CRISPR or linkages.

We next analyzed our timeseries data with CRASS, a bioinformatic tool that re-assembles CRISPR arrays from metagenomic reads to identify additional spacers associated with our host MAGs (58). Briefly, repeat sequences identified within host arrays were matched to those identified with CRASS and spacers grouped to the CRASS-identified repeat were then associated with the MAG. This approach identified thousands of additional spacers from metagenomic reads and allowed us to associate as many of the recovered spacers from all timepoints as possible. Importantly, this facilitated the identification of temporal trends in CRISPR array sizes at both the genome and community level, as insights into host arrays were not limited to a single timepoint where the MAG was recovered.

For many recovered genomes, the number of CRISPR spacers generally exhibited a strong positive relationship with MAG coverage, a proxy for relative abundance in the community (Figure 2 & Figure S8). This trend was obscured when MAGs were grouped together at higher taxonomic levels, highlighting the need for genome-resolved analyses (SI & Figure S9). We also observed a significant positive relationship between the total number of detected viruses and the number of spacers recovered from the metagenome at the community level (Figure S10). Both connections provide further evidence that in natural ecosystems, encoding a CRISPR array and acquiring spacers as a result of viral interactions may support the host’s overall success.

**Figure 2.**
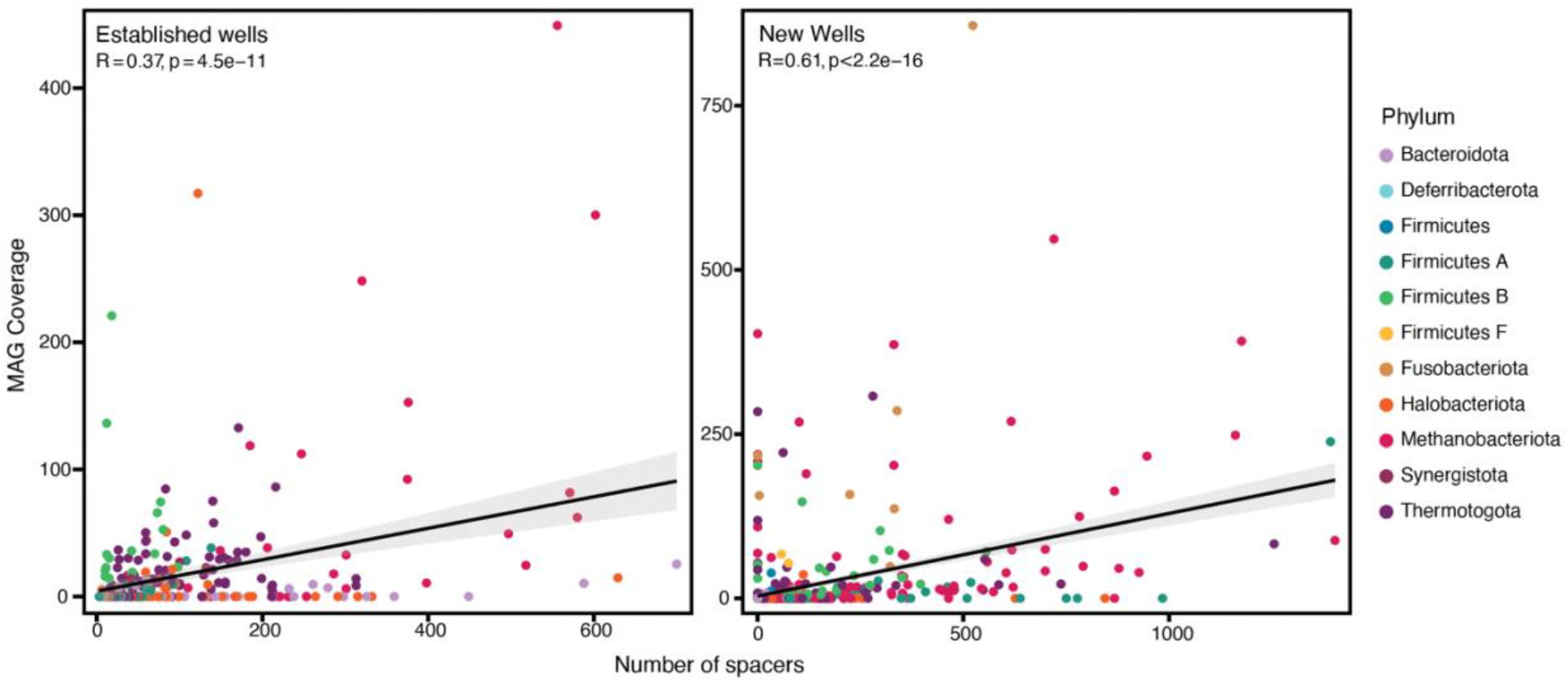
Spearman correlations of host (MAG) coverage and number of spacers associated with each MAG for new and established wells. Each dot represents a single MAG at a single timepoint, colored by phylum.

CRISPR is just one of many viral defense systems, many of which have been only recently described (59). Not all MAGs encoded a detectable CRISPR array, and thus we hypothesized that other viral defense systems are likely deployed by hosts. We found that 87% of all MAGs contained another detectable viral defense system, with no significant difference in the proportion of MAGs between the new and established wells. However, a greater proportion of MAGs lacking CRISPR in the established wells encoded a different viral defense system (82%) compared to the new wells (72%), again highlighting the need to defend against viral predation in this ecosystem and the wide repertoire of systems deployed (SI & Table S2).

### Temporal convergence in patterns of host-virus co-occurrence

Spacers within CRISPR arrays can be uniquely leveraged to make strong inferences about host-virus interactions, as spacers from the host array often identically match the viral protospacer ‘target’ (15). Leveraging additional spacers identified via CRASS, we were able to identify 2,110 viral linkages across 90 MAGs representing 25 different phyla (Figure 1). Indeed, matching all spacers associated with a host MAG to our vMAG database yielded at least one viral linkage for a majority of MAGs encoding a CRISPR array. We observed ≥1 viral linkage for 68% of MAGs with CRISPR arrays in the established wells (48 of 70 MAGs) and 79% of MAGs with CRISPR arrays in the new wells (42 of 53 MAGs) (Figure 1, Table S2). There was no significant relationship between MAG coverage and the number of viral linkages in the established wells, and a weak positive relationship in the new wells (Spearman’s rho *R=0.39, p=1.4e-07*), suggesting that there is not a sustained relationship between the hosts’ overall success and the number of different interacting viruses.

Even amongst widespread CRISPR defense and the presence of matching (linking) spacers, many viruses persisted. Therefore, we next evaluated how often viruses can persist and interact with the host population, despite theoretical CRISPR defense. We leveraged our many host-virus linkages to quantify differences in host-virus co-occurrence patterns in both sets of wells and studied how these dynamics may develop or even equilibrate through time. We set forth three simplified rules defining successful vs. unsuccessful interactions between a virus and host, from the perspective of a virus: (1) when a host was present and the virus was not, the viral interaction was considered unsuccessful, (2) when the virus and host were present, the viral interaction was considered successful, and (3) when the virus was present, but the host was not, the viral interaction was also considered successful (Figure 3).

**Figure 3.**
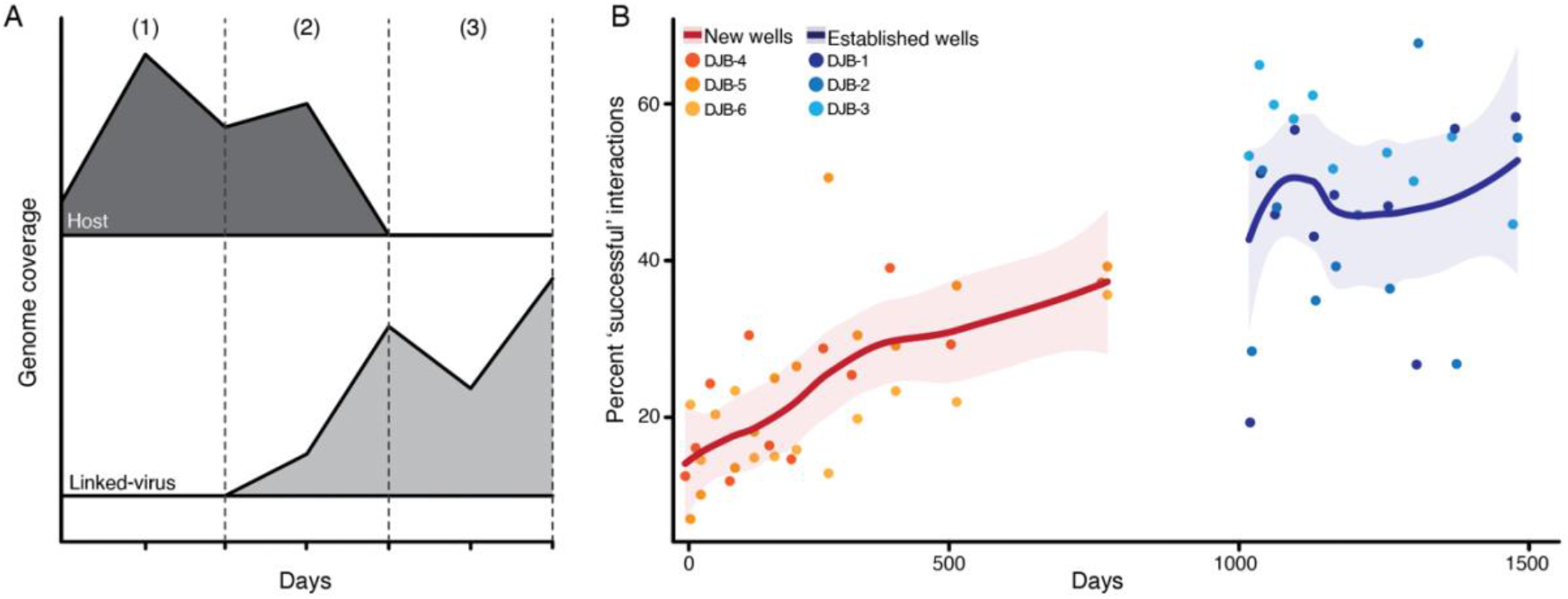
Patterns of host-virus co-occurrences. (A) Conceptual diagram of what is considered a successful vs. unsuccessful interaction. An unsuccessful viral interaction is considered any timepoint where the host was present, but the virus was not present (below detection) (1), while successful viral interactions were considered to be when the virus was present but the host was or was not present (2 and 3). Axes are purposefully left blank given for this conceptual illustration. (B) Temporal trends in percent of successful interactions in the new and established wells.

We observed differing patterns of host-virus dynamics between the new and established wells, potentially reflecting the establishment of more spatially heterogenous microbial communities in the older wells. In the older established wells, we observed an almost even split in occurrences of successful (48%) and unsuccessful (52%) interactions, implying that viruses are generally more present when their hosts are present. In contrast, we observed many fewer occurrences of ‘successful’ interactions (24%) and far more ‘unsuccessful’ (76%) in the new wells, indicating that there are many more instances where a host is present, but the linked virus is absent. Over time however, occurrences of successful/unsuccessful interactions began to shift towards the more balanced split observed in the established wells (Figure 3), suggesting that an equilibrium may develop, resulting in greater host-viral co-existence.

In contrast to expectations that instances of ‘unsuccessful’ viral interactions would increase through time as hosts utilized increasingly effective CRISPR defense systems, we instead observed the opposite trend. We speculate that hosts and viruses in established wells likely exist in more organized spatial structures (i.e., biofilms) that foster continual interactions between the same host and viral populations, in contrast to new wells where less developed biofilms could limit opportunities for stable and continuous host-virus interactions. Only a small portion (<10%) of the recovered vMAGs encoded integrase genes, and very few matched their host coverage, indicating that most viruses are likely lytic and a temperate lifestyle is unlikely to be driving these trends in virus-host co-occurrence (SI). Finally, anti-CRISPR genes were identified in 16 different vMAGs that were more persistent in the established wells. This may be another mechanism behind greater host-virus co-occurrence in the established wells, as viruses encoding these genes may evade CRISPR defenses more successfully. Together, these findings shed light on the complexities of host-virus dynamics both temporally and spatially and how subsurface closed ecosystems may develop towards an equilibrium of host-virus co-occurrences, as opposed to dominance by host or viral populations (60) or ‘red queen’ dynamics of constant evolution and population turnover (61,62).

### CRISPR arrays in the established wells harbor fewer spacers, possibly reflecting development towards optimal size and efficiency

With continued interaction between hosts and viruses, we hypothesized that microorganisms in the established wells would generally encode larger, more expansive CRISPR arrays that reflected these events. However, some studies have noted the loss of spacers through time and others have even proposed a theoretical optimum for CRISPR array size (34–36,42,44). Despite higher host and viral diversity in the established wells, we observed fewer spacers in host arrays recovered in these wells relative to the new wells, indicating that larger arrays are not necessarily driven by higher viral diversity or virus-host interactions. Indeed, arrays from the new wells contained on average a greater number of spacers (avg. 288±410) compared to the established wells (avg. 180±306) (Figures S11 & S12). Given that reductions in immunity have been proposed as a possible driver of host-virus co-occurrence (14,63), the detection of fewer spacers in CRISPR arrays from established wells could account for the co-occurrence patterns observed in Figure 3. Complementing the larger arrays, more viral linkages were also made per MAG from the new wells relative to the established wells, averaging 22 and 7 unique viral linkages, respectively (Figure S13). Less linkages per MAG observed in the established wells may be driven by interactions with fewer different viruses, potentially due to more heterogeneous and confined spatial structures where host and viral populations interact.

Although MAGs from the established wells encoded fewer spacers and linked to less viruses, we observed less redundancy within those viral linkages. For all linkages made (i.e., every host linked to any virus) for MAGs from the established wells, an average of 71% of those linkages were to unique viruses (Figure 4). In the new wells we observed greater redundancy in linkages to the same virus, as only 39% of all linkages were to unique viruses. Therefore, while MAGs from the established wells contained fewer spacers and linked to less viruses, they could be matched to a proportionally greater number of different viruses, suggesting that the retained spacers protect against a wider suite of viruses with less redundancy.

**Figure 4.**
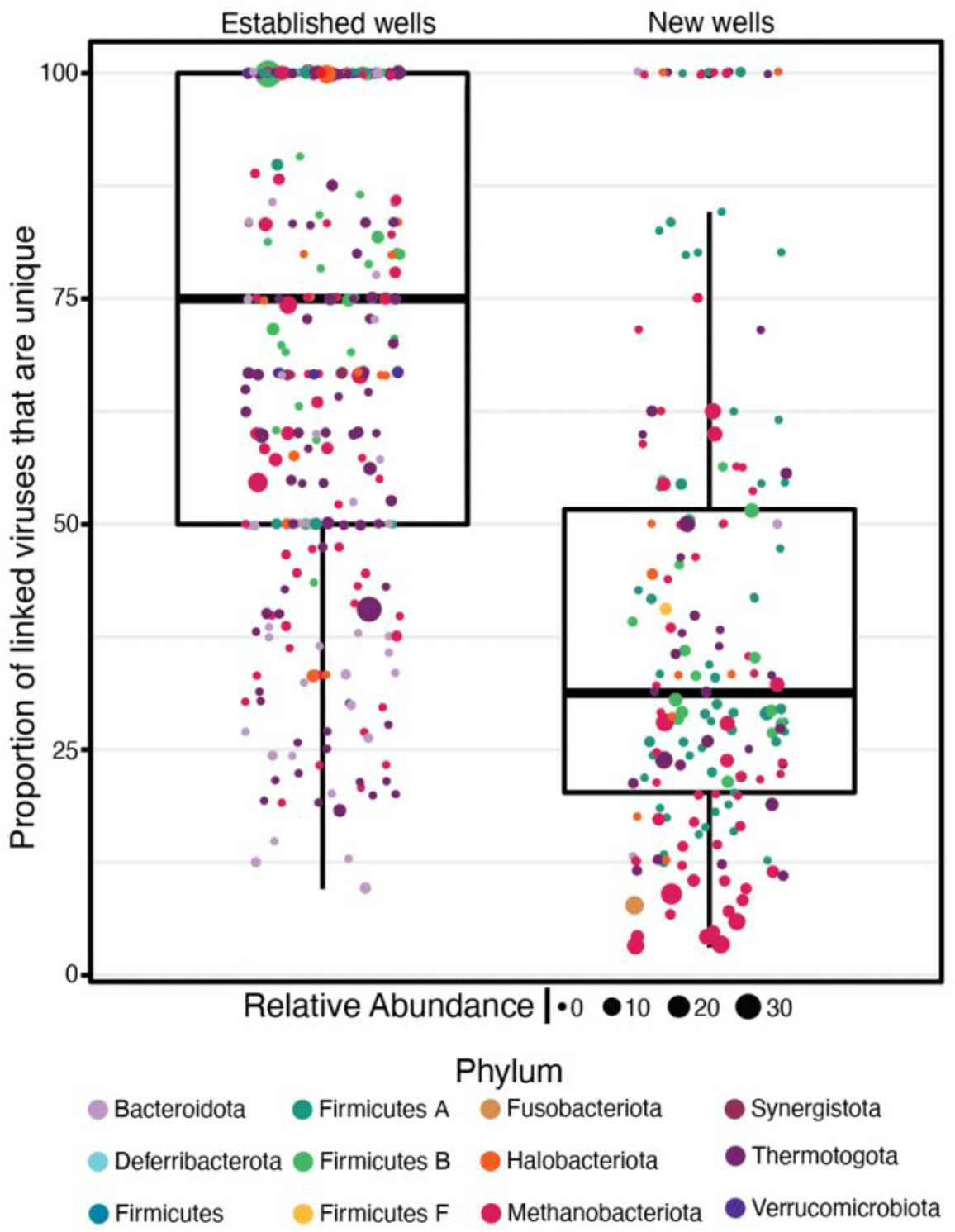
Boxplots illustrating the proportion of linked viruses, per MAG, that are unique. Each point represents a single MAG at a single timepoint where spacers were recovered and linkages are made, colored by phylum and sized by relative abundance. The proportion of unique viruses linked was calculated as the number of different viruses linked to out of the total number of linkages at a given timepoint.

There are myriad factors that may limit the size of CRISPR arrays in any ecosystem. Smaller arrays may even be favored, providing the array still confers sufficient defense against a wide enough repertoire of viruses that the host may encounter (40,42,44). For example, larger arrays have been shown to have a greater chance of self-targeting, and thus smaller arrays may be necessary to avoid host autoimmunity (64). We propose that the difference in size of arrays between hosts in the two sets of wells may reflect a temporal shift towards a theoretical optimum in array size that has been suggested through modeling studies (roughly a dozen to one hundred spacers in natural communities) (42,44). In this framework, the optimal efficiency of CRISPR defense may rely on maintaining select spacers rather than retaining all memories of host-virus interactions. Spacers that are retained may match protospacers with fewer mutations or perhaps evolutionarily conserved regions of the viral genome. Thus, under viral predation, Cas machinery is most efficient when not overwhelmed by excessive spacers and deployed with arrays containing less ‘dilution’ effects by older, and possibly less effective, spacers (8,41). As such, there is likely a benefit to forgetting interactions and maintaining smaller arrays, such as those we report in MAGs recovered from the older, established wells.

A smaller array may be more effective against viral predation if the retained spacers target regions of the viral genome with fewer mutations. Here, we used single nucleotide polymorphism (SNP) frequency as a proxy for sequence variation in viral genes to investigate spacer effectiveness. We observed that spacers recovered at only one timepoint generally matched viral protospacers with the highest SNP frequency (Figure 5). Additionally, the percentage of viral genes with zero SNPs increased with spacer persistence in the new wells, but not the established wells (Figure 5). That is, spacers that were present across the most timepoints tended to target virus protospacer genes that had little to no sequence variation within the community. In general, more spacers from the established wells persisted for at least half the sampling timepoints compared to the new wells, with 4.5% and 1.3% of spacers present in at least half the timepoints for the established and new wells, respectively. These trends are likely associated with the increased selection and retention of spacers that most likely confer viral resistance for longer periods of time, and that subsequently allow movement towards an optimal array size in these natural ecosystems.

**Figure 5.**
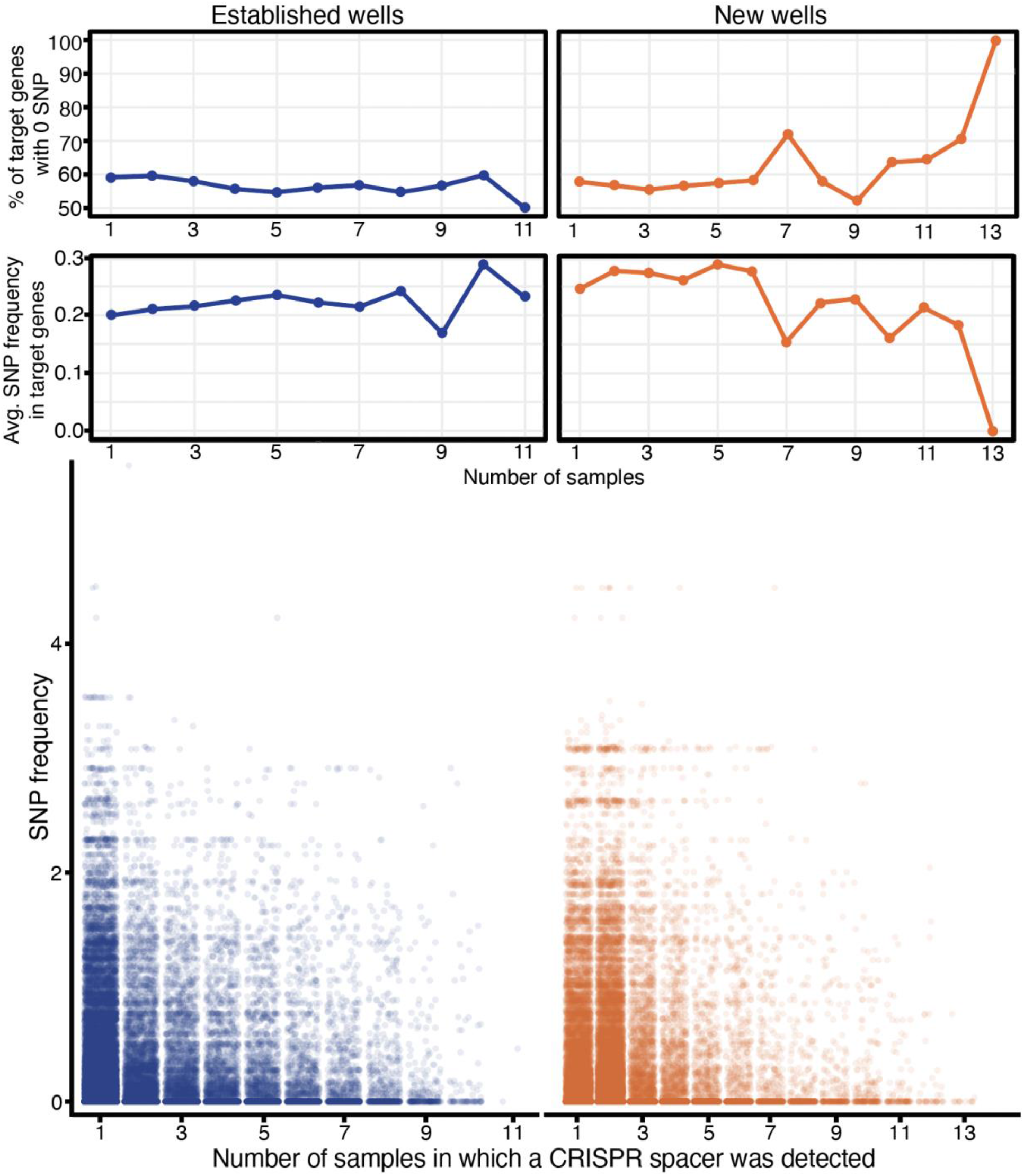
SNP frequency (number of SNPs/gene length) for protospacer genes with corresponding persistence of the host spacer. Spacer persistence is quantified as the number of samples that the spacer was recovered. Average SNP frequency as well as percent of viral genes with zero SNPs is shown as line graphs above raw data plots.

Fewer spacers within an array may also help to reduce the possible dilution effect caused by excess spacers paired with Cas machinery (8,40). Upon activation of CRISPR defense, spacers are transcribed and partner with Cas nucleases to form effector complexes that target and degrade invading viral nucleic acids. Extensive array sizes can reduce the efficacy of the deployed Cas machinery, as most recently encoded (and most likely to be effective) spacers are ‘diluted’ by older and likely less effective spacers. Hosts from the new wells, which generally had a greater number of spacers associated with them, also tended to encode more distinct arrays compared to the established wells. Having multiple distinct arrays is likely key in supporting large numbers of spacers that are distributed amongst several arrays instead of just one. Even so, array sizes in the new wells were generally larger than those in the established wells. Thus, smaller arrays may be favored if the effector complexes can concentrate on a smaller pool of spacers, rather than mounting a viral defense with similar numbers of Cas genes but hundreds of spacers.

## Conclusion

Here we leveraged time-resolved samples from two sets of hydraulically fractured shale wells to establish CRISPR-based host-viral linkages and study long-term host-virus co-existence and CRISPR array dynamics. Data from six individual timeseries (>500 days) allowed us to recover CRISPR repeats and spacers from metagenomes and MAGs and identify host-virus linkages and temporal dynamics of CRISPR array sizes. CRISPR viral defense systems were widely encoded with CRISPR arrays detected in the majority of MAGs, together representing 25 of 29 different phyla within the microbial communities. Viral linkages were made for a majority of hosts containing a CRISPR array, with over two thousand total linkages identified across 90 different MAGs. Given the prevalence of CRISPR defense systems and the important role this defense might have in host-viral co-existence, we interrogated host-viral temporal dynamics and found that co-occurrence of host and viral populations generally increased through time, potentially due to the development of more spatially heterogeneous microenvironments (i.e., biofilms) in the deep subsurface. Finally, we demonstrated that CRISPR array sizes in the older wells were generally smaller than those recovered from the new wells and that spacers from the old wells could potentially provide longer-term defense through targeting viral sequences with less nucleotide variation. This provides additional evidence that hosts in natural communities may preferentially maintain smaller and more efficient arrays rather than retaining all memories of past viral infections. Together, this study offers new insights into the long-term dynamics between host and viral populations and CRISPR-based host-viral dynamics within a natural ecosystem.

## Materials & Methods

### Recovery and processing of samples from the DJ Basin

Produced fluid samples were collected from six hydraulically fractured wells from the Niobrara formation, within the Denver-Julesburg (DJ) Basin, in eastern Colorado between October 2018 and October 2020 (*n=78*) (Figure S1). Additional details on the hydraulic fracturing of these subsurface shales are described in the SI. The six wells are split equally into two groups defined by their age when sample collected began: the three ‘established’ wells (*n=33*) had been producing for approximately 1000 days prior to sample collection (DJB-1, DJB-2, and DJB-3), while the three ‘new’ wells (*n=45*) were sampled from day ~30 in production (DJB-4, DJB-5, and DJB-6). A small number of early produced fluid samples (those beginning with ‘JMDJ#’, <60 days) were collected directly from well heads and filtered through a 0.22μm pore size polyethersulfone membrane Sterivex filter (MilliporeSigma) with a minimum of 500mL of fluid filtered. Most produced fluids (those beginning with ‘DJKA’) were collected directly from separator tanks into 1L Nalgene bottles with no head space and stored at 4°C until processing, which occurred within 24 hours from when the sample was collected. Between 500-800mL of fluid was filtered through a 0.2μm PES membrane Nalgene vacuum filtration unit (Thermo Scientific). Filters were removed from the units and stored at −20°C until DNA extraction. Conductivity was measured on raw, unfiltered fluids at room temperature using a Myron L 6PIIFCE meter.

### DNA extraction and metagenomic sequencing

Total nucleic acids were extracted from half of each sample’s 0.2μm filter using DNAeasy PowerSoil Kit (Qiagen). Extraction blanks were run with each round of DNA extractions and all returned no detectable nucleic acids using the maximum amount of blank sample (20μL) via the Qubit dsDNA High Sensitivity assay kit (ThermoFisher Scientific). For all 78 samples, genomic DNA was prepared for metagenomic sequencing at the Genomics and Microarray Core at the University of Colorado, Denver’s Genomics Shared Resource. Samples were prepared using the Illumina Nextera XT Library System according to manufacturer’s instructions for 2×150bp libraries and were sequenced using the Illumina NovaSeq platform and paired-end reads were collected.

### 16S rRNA gene sequencing and analysis

Nucleic acids for all samples were also sent to Argonne National Laboratory for 16S rRNA gene sequencing (Table S1). Sequencing was performed with the Illumina MiSeq platform, using the Earth Microbiome Project primer set for amplification of the 251bp hyper-variable V4 region. 16S rRNA gene sequences were obtained via Argonne’s standard procedure, with the exception of performing 30 PCR amplification cycles. Paired-end reads were processed with QIIME2 (v 2021.2) EMP protocol, by first demultiplexing via exact-matching of barcodes, trimmed to 250bp and denoised with DADA2 (65). Representative sequences were taxonomically classified with SILVA (release 138). All 16S rRNA gene sequencing reads were submitted to NCBI under BioProject PRJNA308326 and individual accession numbers are listed in Table S1.

### Metagenomic assembly, binning, and viral recovery

For bacterial, archaeal, and viral recovery, total sequenced DNA from each sample was first trimmed from 5’ to 3’ ends with Sickle (https://github.com/najoshi/sickle) and individually assembled using IDBA-UD with default parameters (66). Assembly information for each sample is provided in Table S1. Only scaffolds ≥5kb from metagenomic assemblies were used for binning bacterial and archaeal genomes with MetaBAT2 (v2.12.1) to recover metagenome assembled genomes (MAGs) (67). CheckM (v.1.1.2) lineage workflow (‘lineage_wf’) followed by the ‘qa’ command was used to assess completion and contamination for each metagenomic bin (68), and medium (>50% completion, <10% contamination) and high (>90% completion, <5% contamination) quality bins were recovered from all samples from all six wells following the standard metrics for MAGs proposed by Bowers et al. (69). The two sets of unique MAGs (from the new and established wells) were individually determined by dRep v2.2.3 using default parameters (70). We refer to the final set of 202 MAGs as the ‘host’ community. All MAGs were taxonomically classified using GTDB-Tk v2.2.0 (71). Metagenomic assemblies and MAGs were annotated via DRAM (v1.2.4) using default parameters (72). Additional details about MAGs can be found in Table S2.

Viral MAGs (vMAGs) were also identified in metagenomic assemblies from scaffolds ≥10kb in length using VirSorter2 (v2.2.2) (73) and following the “Viral sequence identification SOP with VirSorter2” developed by the Sullivan Lab (74). Following this protocol, quality of vMAGs were assessed using checkV (v0.8.1) and annotated using DRAM-v (v1.2.4) (72,75). Low confidence vMAGs were removed following the manual curation steps in the SOP. Viral genomic contigs (≥10kb) were clustered into viral populations (genus level) using the ‘ClusterGenomes’ (v 1.1.3) app in CyVerse using the parameters 95% average nucleotide identity and 90% alignment fraction of the smallest contig (https://github.com/simroux/ClusterGenomes). The resulting database of 2,176 vMAGs are considered our viral database. Viral taxonomy was determined by clustering vMAGs with viruses belonging to the viral reference taxonomy databases in NCBI Bacterial and Archaeal Viral RefSeq v211, and viruses from the International Committee on Taxonomy of Viruses (ICTV) via vConTACT2 v0.11.3 with default settings (76). Anti-CRISPR (arc) genes were identified in vMAGs using ArcFinder (using both homology-based and guilt by association based approaches) with default parameters (77). Additional details about vMAGs can be found in Table S3.

### Calculating MAG and vMAG coverage and relative abundance

To calculate coverage and relative abundance of MAGs and vMAGs, all 78 pairs of trimmed metagenomics reads were rarified to the lowest metagenome sequencing depth of 9Gbp using the ‘reformat’ guide within bbmap (78). Coverage for MAGs was calculated by competitively mapping rarified metagenomic reads to MAGs using bbmap (v38.89) with minid=90. Resulting sam files were converted to sorted bam files using samtools (v1.9) (79). Coverage for each MAG was calculated using coverM (v0.6.0) (https://github.com/wwood/CoverM) using two commands. First, coverM was run using –min-covered-fraction=90 to determine MAGs read recruitment to at least 90% of the genome. Second, coverage values were calculated using the -m reads_per_base command, which represents reads mapped/genome length, and thus multiplied this by read length (151bp) in order to calculate MAG coverage (simply, coverage = reads_per_base * 151 bp). Only MAGs with >1x coverage and with reads mapped to >90% of the genome were considered present in a sample. Relative abundance was thus calculated as the proportion of a given MAG’s coverage out of the sum of all present MAGs’ coverage, per sample.

Metagenomic reads were also mapped to vMAGs to determine coverage using bbmap with minid=95 (v38.89)(78) and sam files converted to bam files using samtools (v1.9) (79). Given vMAGs are viral contigs, coverM (https://github.com/wwood/CoverM) contig mode was applied with two commands. First, --min-covered-fraction 75 and next followed by -m reads_per_base to calculate coverage. Similar to requirements set for MAGs, here vMAGs must have a minimum covered fraction >75% to be considered present. Coverage values were calculated from the reads per base output*151 bp. Number of viruses present in a metagenome were determined by presence of vMAGs given this recruitment of metagenomic reads.

### Detection of viral defense systems and recovery of spacers

CRISPR arrays in MAGs were identified using the Geneious (v.2020.0.5) plugin CRISPR Recognition Tool (CRT) (80) v.1.2 using the ‘Find CRISPR loci’ annotation tool with the following parameters: min number of repeats a CRISPR must contain: 4, minimum length of a CRISPR’s repeated region: 19, maximum length of a CRISPR’s repeated region: 55, minimum length of a CRISPR’s non-repeated region (or spacer region): 19, maximum length of a CRISPR’s non-repeated region (or spacer region), length of a search window used to discover CRISPR’s: 8. Spacers were also detected in non-rarified and rarified trimmed metagenomics reads using CRisprASSembler: CRASS (v1.0.1) (58). Briefly, CRASS reassembles CRISPR-Cas arrays of repeats and spacers that tend to break during assemblies and groups spacers by the repeat sequences in CRISPR arrays. Only the total of spacers recovered from rarified metagenomics reads were used to represent the ‘total number of spacers in a metagenome’ for all community-level correlations to not introduce bias from varying read depth into these analyses. All recovered host genomes, regardless of detection of a CRISPR array, were also queried for 60 other known anti-phage systems using DefenseFinder (v.1.0) (59).

### Making CRISPR-based host-virus linkages

Linkages between MAGs and vMAGs (hosts and viruses) were made exclusively via CRISPR spacers using two approaches. As a result of this, linkages could only be made with MAGs that had a detectable CRISPR array. First, CRISPR arrays were identified in MAGs using Geneious, and spacers and repeats were extracted from the CRISPR arrays. We then leveraged CRASS to make as many linkages as possible and evaluate the number of spacers associated with a MAG through time. Repeat sequences from MAGs were identically matched to direct repeat sequences from CRASS (same length, no mismatches). Spacers that were associated with a direct repeat sequence from CRASS were thus grouped with the MAG of the same repeat sequence. To make as many host-viral linkages as possible, spacers were extracted from CRASS applied to non-rarified reads. Next, spacers from all MAGs (linked via Geneious and CRASS) were queried against all vMAGs using BLASTn with the parameters to optimize short sequences BLAST: -dust no and -word_size 7. Finally, only identical or nearly identical (0 or 1 mismatch across spacer length) were used to match spacers to vMAGs and make host-viral linkages.

### Host-viral co-occurrence patterns

All MAGs with at least one viral linkage were included in analyzing host-viral co-occurrence patterns. For each MAG and individual linked virus at every timepoint, all possibilities were evaluated for being ‘successful’ or ‘unsuccessful’ viral interactions. Successful interactions are considered any instance where the virus is present and the host is either present or absent (below detection). Instances where both host and virus were not present were excluded from any calculations and not counted in the total number of interaction occurrences. Thus, the percent of successful viral interactions was calculated as the proportion successful out of all interactions that met the successful/unsuccessful criteria. All early samples that required field filtration (“JMDJ#”) from the new (*n=9*) and established wells (*n=3*) were removed from the temporal graph (Figure 3), as difference in filtering methodology likely influenced this analysis.

### Analysis of single nucleotide polymorphisms in vMAGs

We combined SNP values for viral genes with the persistence of spacers that link host and virus to interrogate any possible relationship between gene variation and spacer retention for all MAGs with linkages. We utilized MetaPop (81) with default parameters to calculate the number of SNPs within all viral genes identified in our vMAGs. For genes that met MetaPop’s default parameters, SNP frequency was calculated relative to the gene length. Genes containing linked protospacers that did not meet MetaPop’s default parameters were not included in this analysis. Finally, SNP frequency for the gene containing the protospacer was combined with the persistence of the spacer (i.e., number of samples the spacer was recovered).

### Statistical analyses

Alpha diversity (Shannon’s index) and beta diversity (Bray-Curtis) values were calculated using vegan v2.6-2 in R. Alpha diversity was calculated using 16S rRNA amplicon data, while beta diversity and Bray-Curtis dissimilarity values were calculated based on the host and viral communities recovered via metagenomic sequencing and rarified reads. Metagenomics was used here since the viral community was recovered using metagenomics and thus the paired host communities were assessed similarly via MAGs (recovered from metagenomes). Bray-Curtis dissimilarity values were calculated as the difference in beta diversity from the previous timepoint. Spearman correlations and p-values were calculated using ggpubr to determine the strength and directionality of relationships between variables such as number of spacers, MAG/vMAG coverage, time, etc. Specifically, correlations between number of spacers and host coverage were only calculated for MAGs that were both present in at least 3 timepoints and also had spacer recovery from at least three timepoints.

## Acknowledgements

This work was supported through an award (EAR-1847684) from the National Science Foundation Geobiology and Low Temperature Geochemistry program to MJW. The work conducted by the U.S. Department of Energy Joint Genome Institute (https://ror.org/04xm1d337), a DOE Office of Science User Facility, is supported by the Office of Science of the U.S. Department of Energy operated under Contract No. DE-AC02-05CH11231 (SR).

