## Supplementary Information for "Long-term CRISPR array dynamics and stable host-virus co-existence in subsurface fractured shales"

### Supplementary Tables

Table S1: Metagenome sequencing and assembly details & 16S rRNA amplicon sequencing details

Table S2: Additional details, including other viral defense systems detected, for all 202 MAGs

Table S3: Additional details for all 2,176 vMAGs, host-viral linkages for MAGs and vMAGs in the new and established wells

### Supplementary Figures

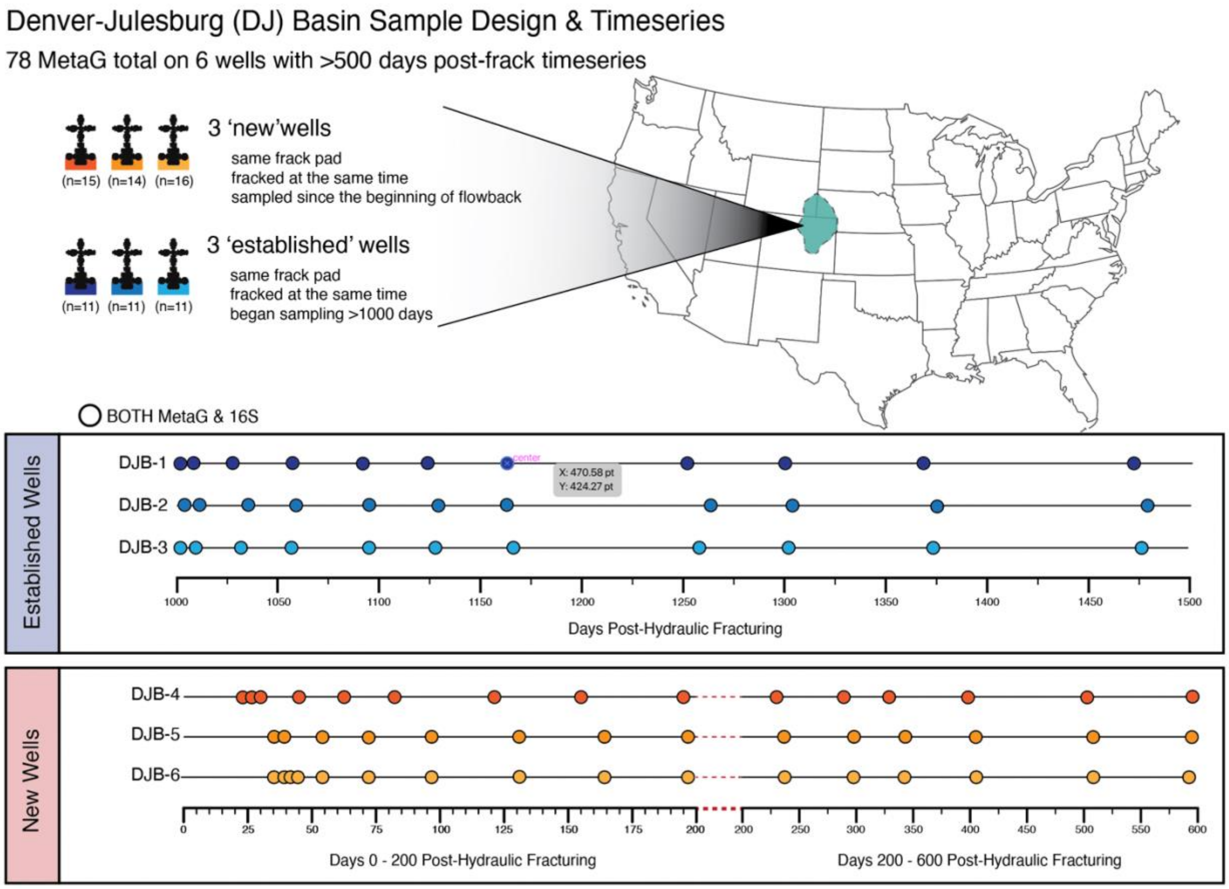


**Figure S1.** Sampling campaign for the six hydraulically fractured shale wells within the Denver-Julesburg (DJ) Basin (Niobrara formation). An approximate outline of the DJ Basin is shown in light blue on the United State map. The six wells were split equally into two groups of three based on well age, relative to when the wells were developed and hydraulically fractured. Metagenomic and 16S sequencing was performed on all samples.

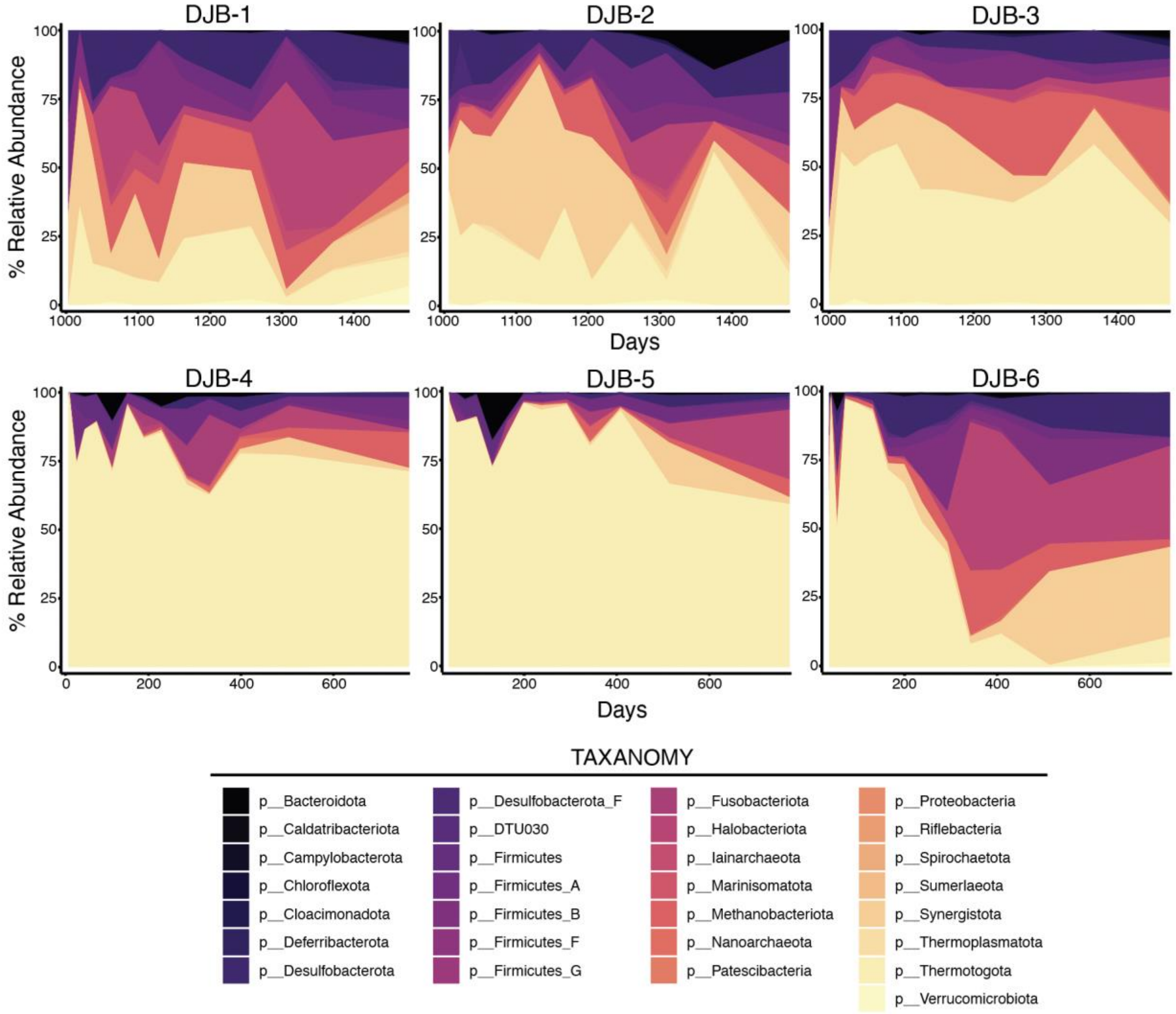


**Figure S2.** Temporal dynamics of the microbial community as shown by MAGs relative abundance in all six wells, summed and colored at the phyla level. Established wells are shown in the top three panels (DJB-1, 2, & 3), while new well dynamics are shown as the bottom panels (DJB-4, 5 & 6).

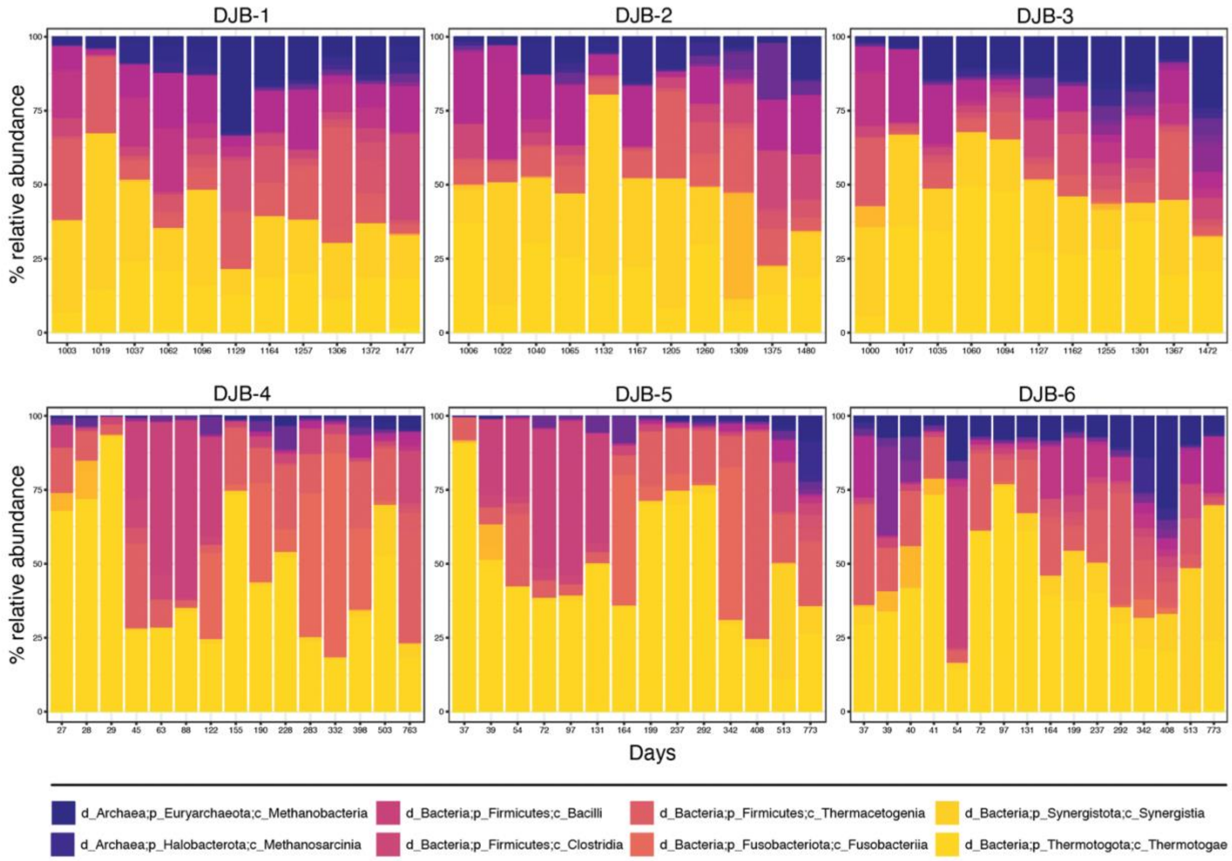


**Figure S3.** Barcharts illustrating the temporal dynamics of of DJ Basin microbial communities based on16S rRNA gene amplicon data. Select dominant taxa that were observed in 3+ wells are highlighted below bar charts.

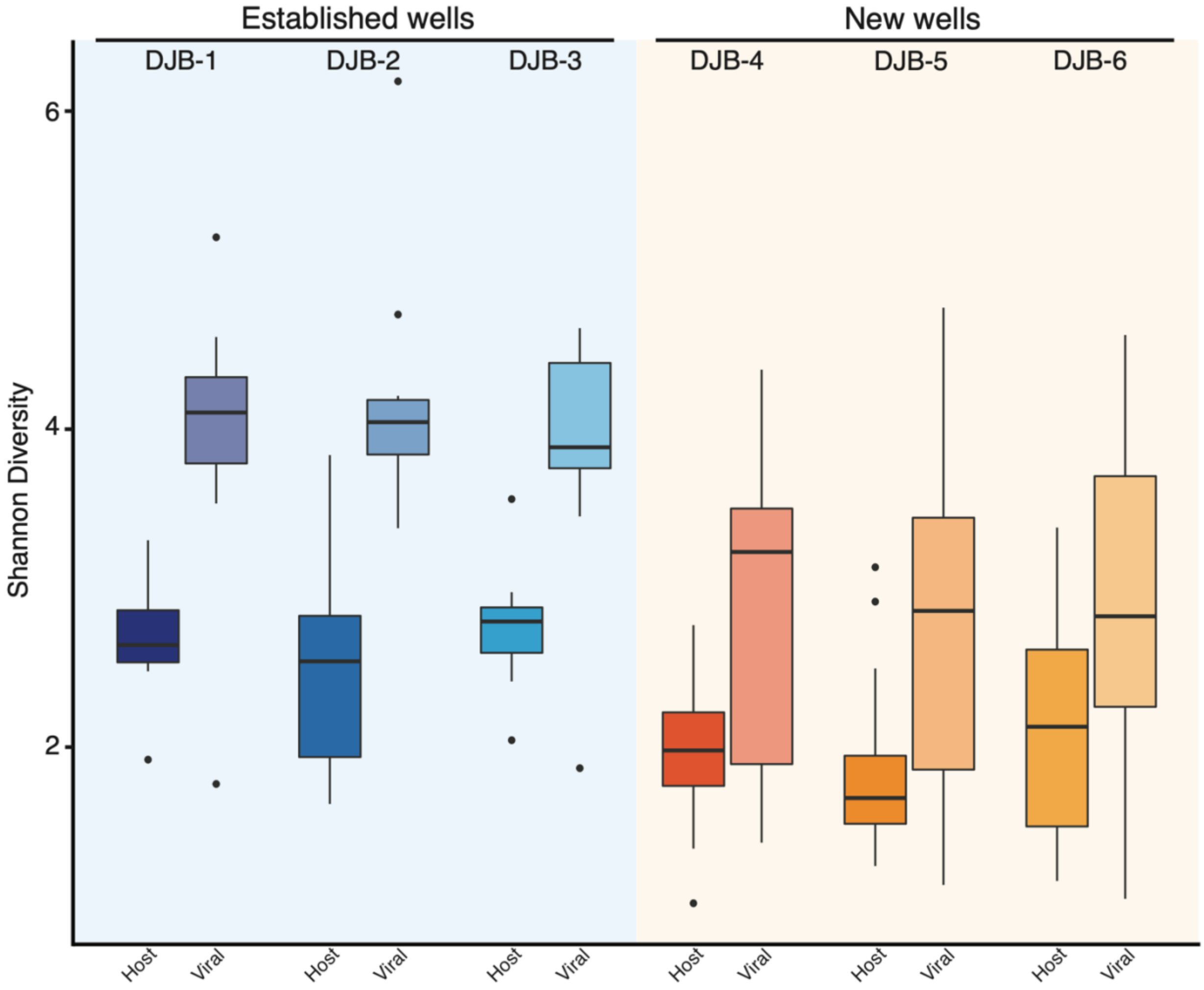


**Figure S4.** Alpha diversity (Shannon's) for both host and viral diversity for all six wells. Host alpha diversity is calculated from 16S rRNA gene amplicon data, while viral diversity was, by necessity, calculated the relative abundance of viral MAGs (vMAGs) – as calculated by metagenomic read recruitment to the dataset of 2,176 vMAGs.

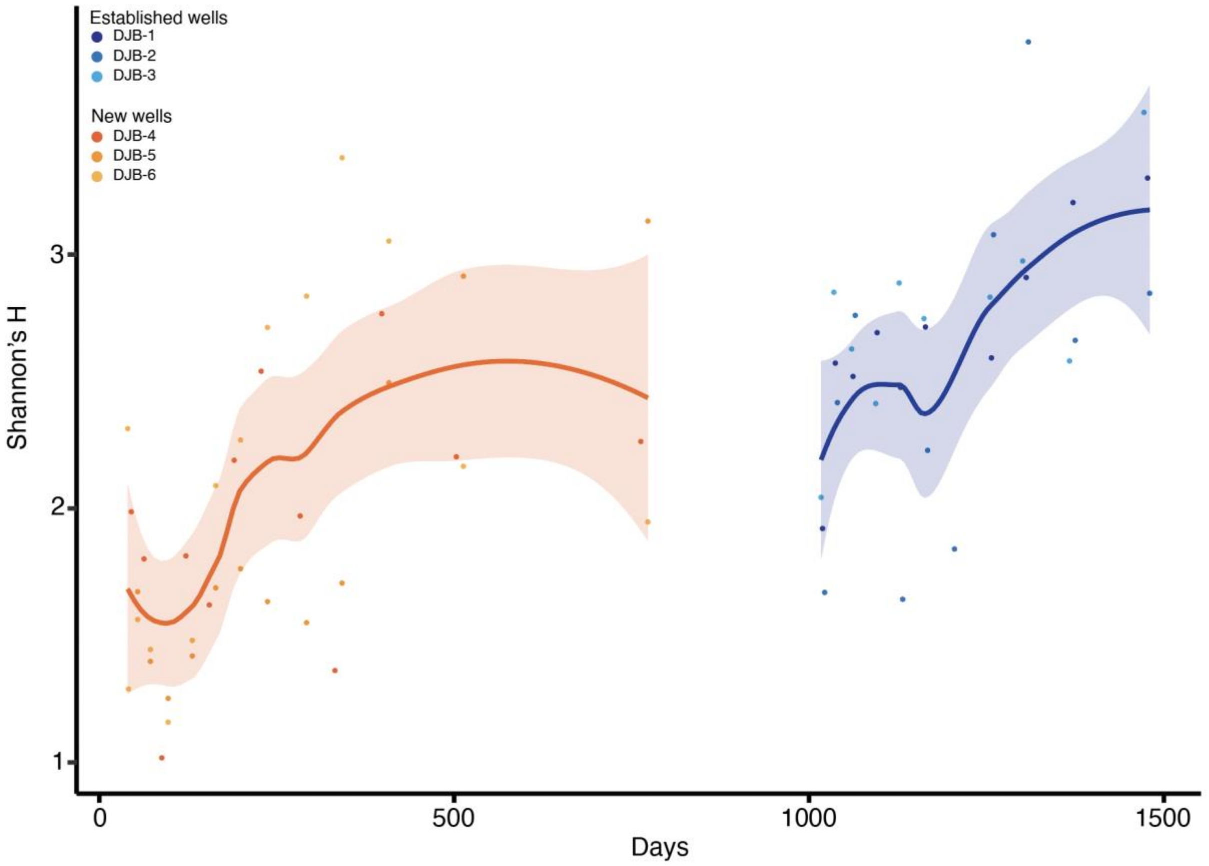


**Figure S5.** Alpha diversity of 16S rRNA gene amplicon data (Shannons') through time reveals a temporal increase in bacterial & archaeal alpha diversity across all six wells.

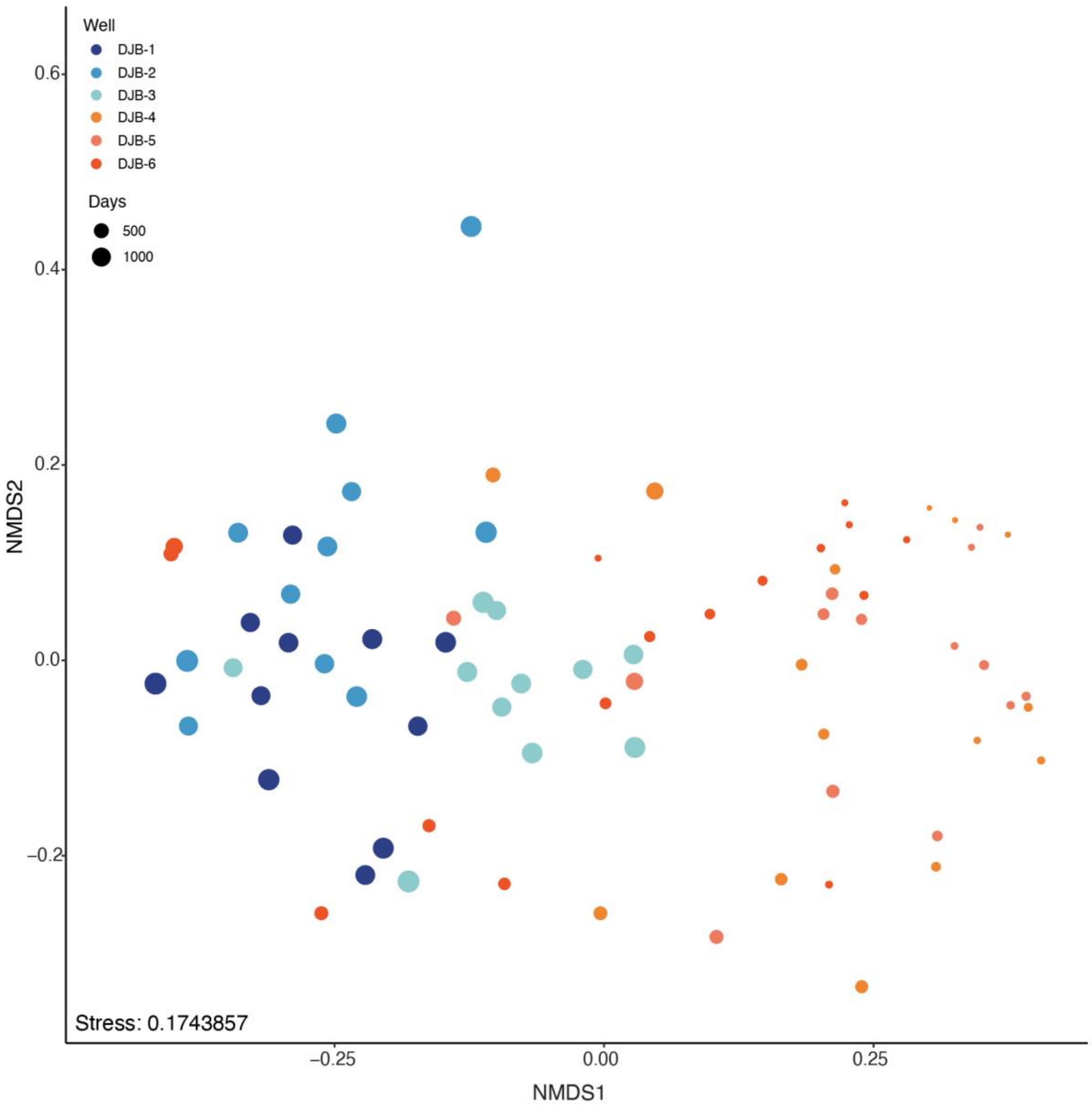


**Figure S6.** NMDS ordination of complimenting 16S rRNA gene amplicon data for the six hydraulically fractured shale wells in the DJ Basin. Points are colored based on well and frackpad (oranges: new wells, blues: established wells) and sized based days since hydraulic fracturing. Samples cluster significantly based on well grouping, however microbial communities from the new wells appear to become more similar to the established well samples through time.

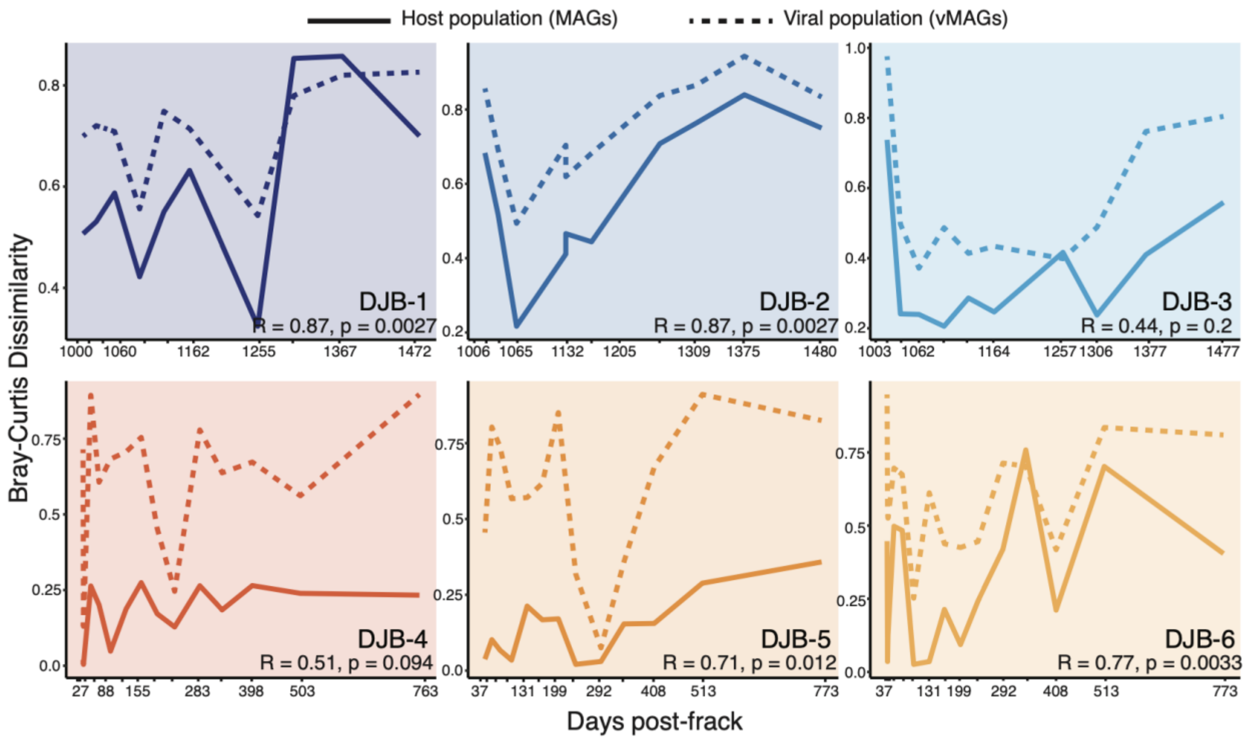


**Figure S7.** Temporal dynamics of host (bacterial and archaeal MAGs) and viral (vMAGs) communities. Bray-Curtis community dissimilarity through time for both host (solid line) and viral (dashed line) communities illustrating the change in each community composition from the previous timepoint, with larger dissimilarity values indicating greater change in community structure. Spearman's rho and p values highlight the strong positive relationship between the temporal changes in host and viral populations in all wells.

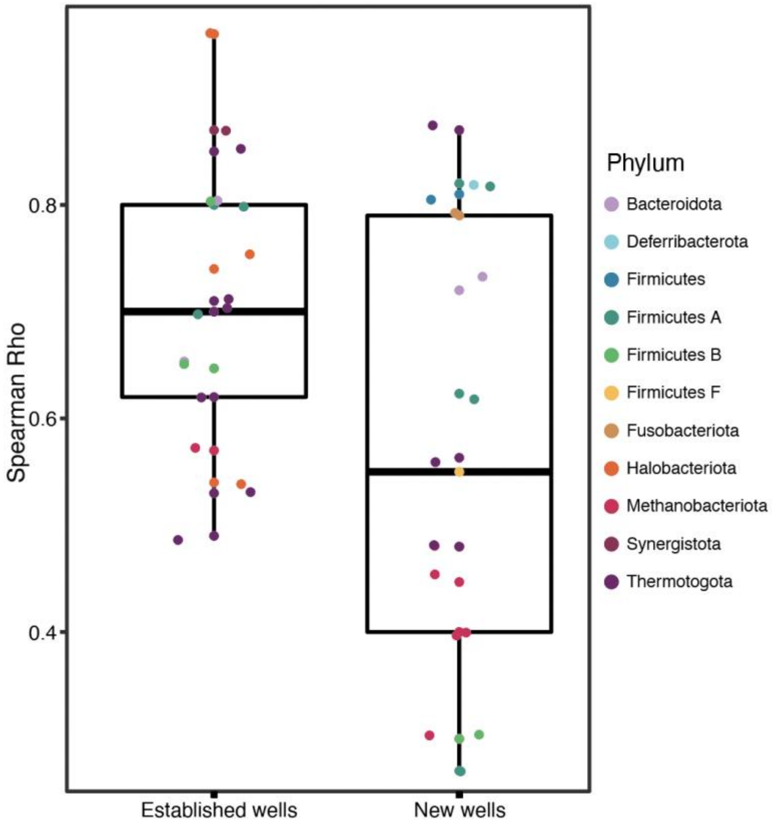


**Figure S8.** Boxplots illustrating genome-level spearman correlations between host (MAG) coverage and number of spacers associated with a MAG. Each point represents the rho value for one MAG (colored by phylum).

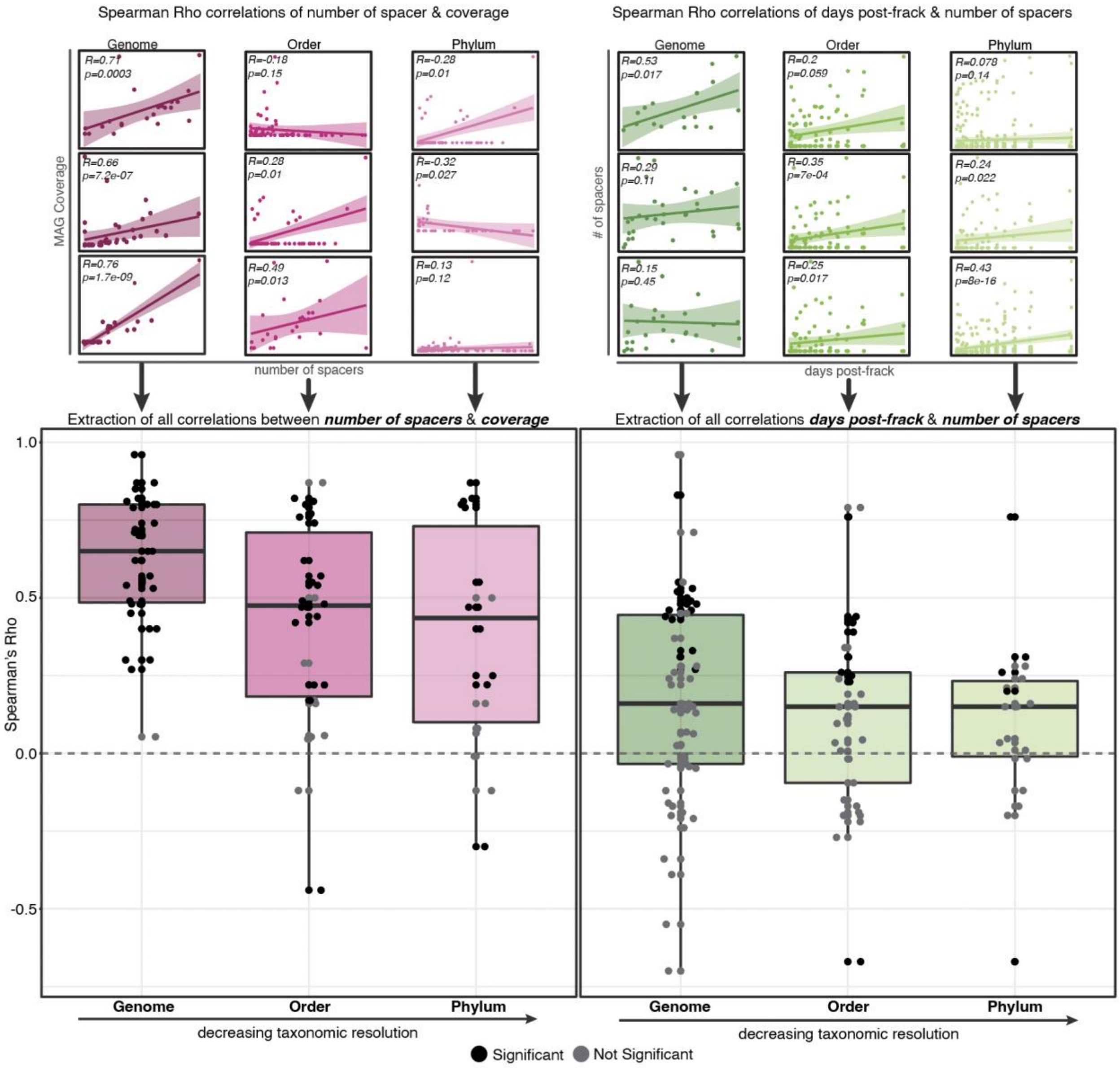


**Figure S9.** Correlations between number of spacers associated with a MAG and MAG coverage or days post frack at three decreasing levels of taxonomic resolution. Top panels illustrate the generation of data (Spearman's rho) that is represented in the boxplots below, with three examples provided per taxonomic level. All rho values and p values for each individual genome, order, or phyla were extracted and plotted as an individual data in the boxplots below for both variables: coverage (pink) or days post-frack (green). Non-significant (>0.05) correlation values (grey) are shown to highlight the lack of a significant positive or negative relationship in many cases, especially at higher taxonomic levels, which is also highlighted in examples above.

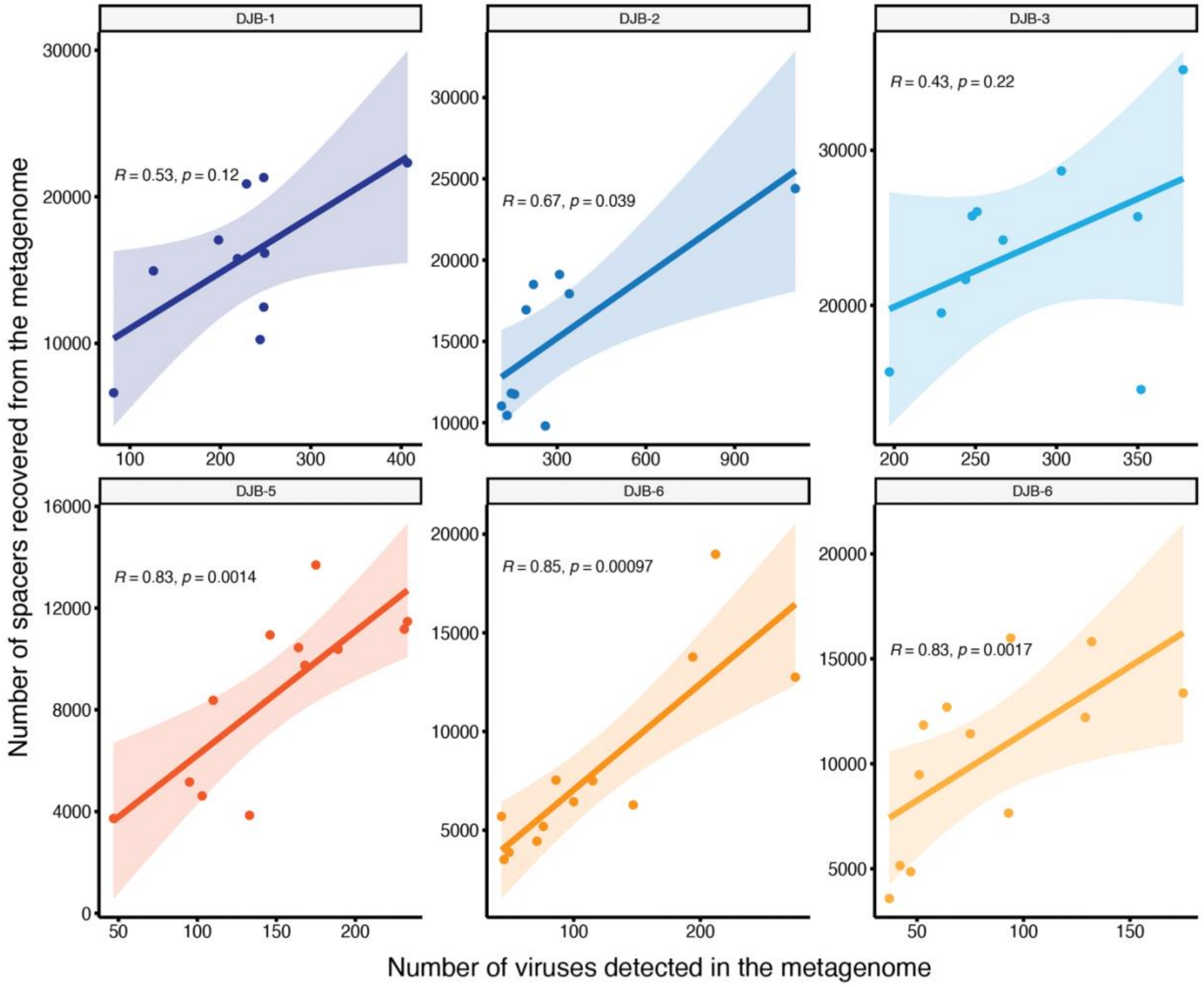


**Figure S10.** Spearman correlations between the number of viruses detected in a metagenome and number of spacers recovered, highlighting a strong positive relationship between them in all six wells. Number of viruses was calculated using mapping of rarified reads to all 2,176 recovered vMAGs. Number of spacers was quantified per metagenome based on spacers recovered via CRASS from rarified metagenomic reads.

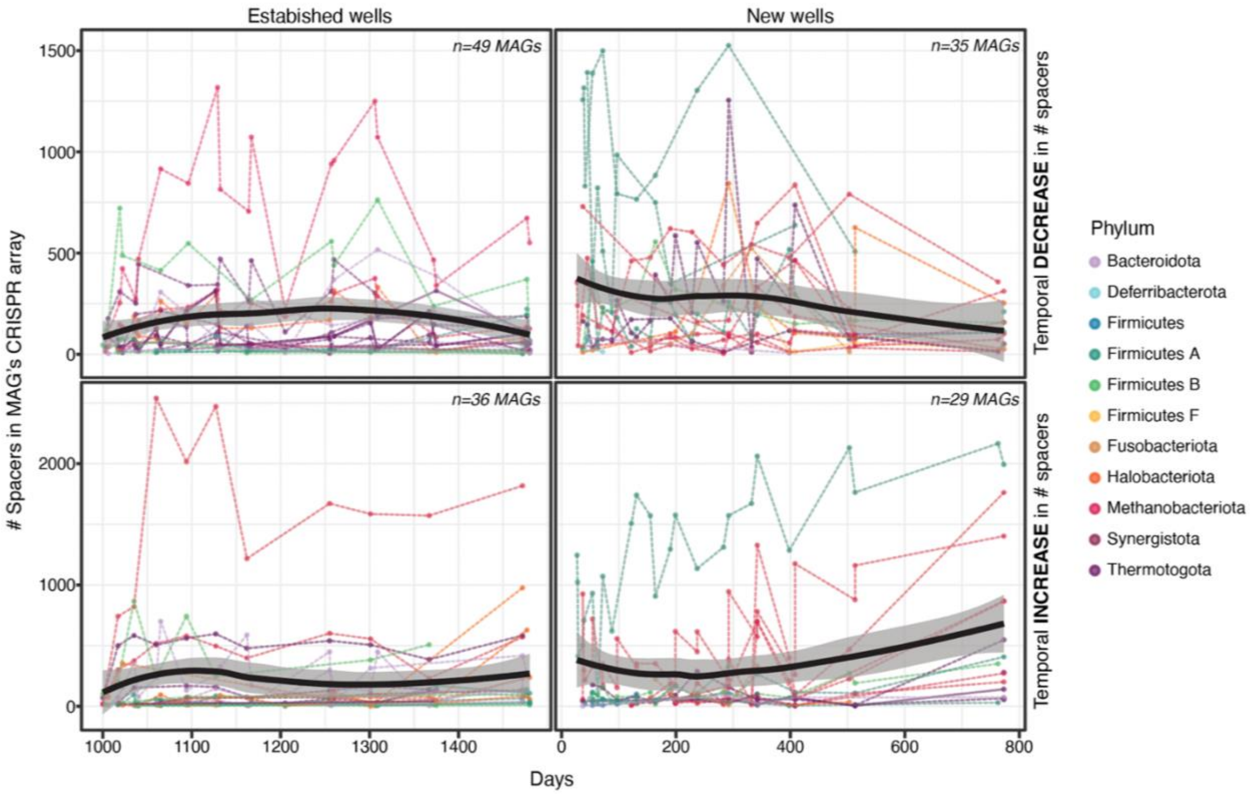


**Figure S11.** Temporal changes in the number of spacers in a MAG CRISPR array. Each dotted line represents a single MAG with general trends highlighted as solid black lines. Top panels show MAGs with arrays that generally decreased (final timepoint < average array size) for the established (left) and new (right) wells, while bottome panels show MAGs with arrays that generally increased (final timepoint > average array size). In general, MAG arrays were smaller and fluctuated less in the established wells than in the new wells.

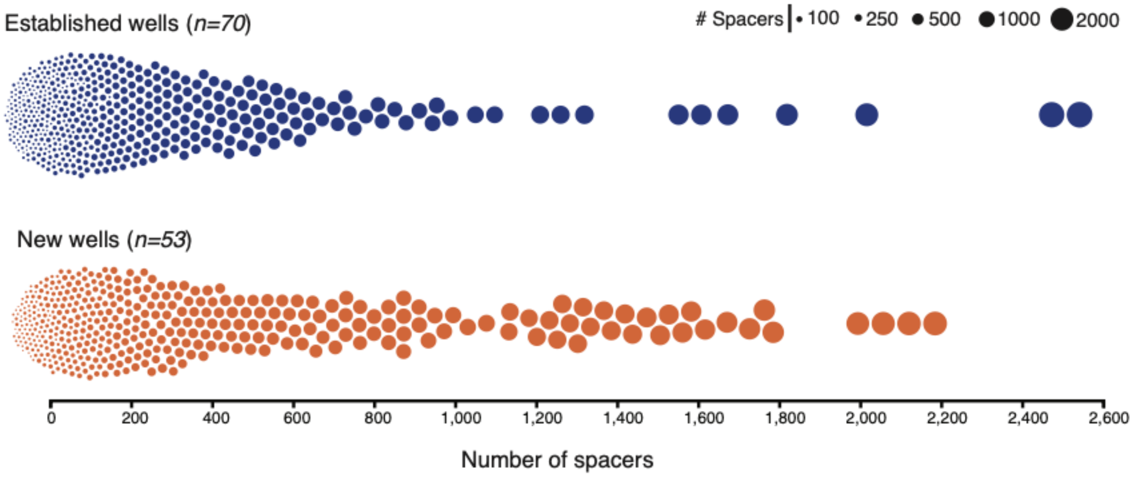

**Figure S12.** Range in the number of spacers in MAG CRISPR arrays. Each dot is both sized and arranged by the size of the CRISPR arrays, colored by new or established wells. Additionally, each dot represents one MAG at one timepoint, and therefore may be multiple points for MAGs that had spacers recovered at alternate timepoints via CRASS.

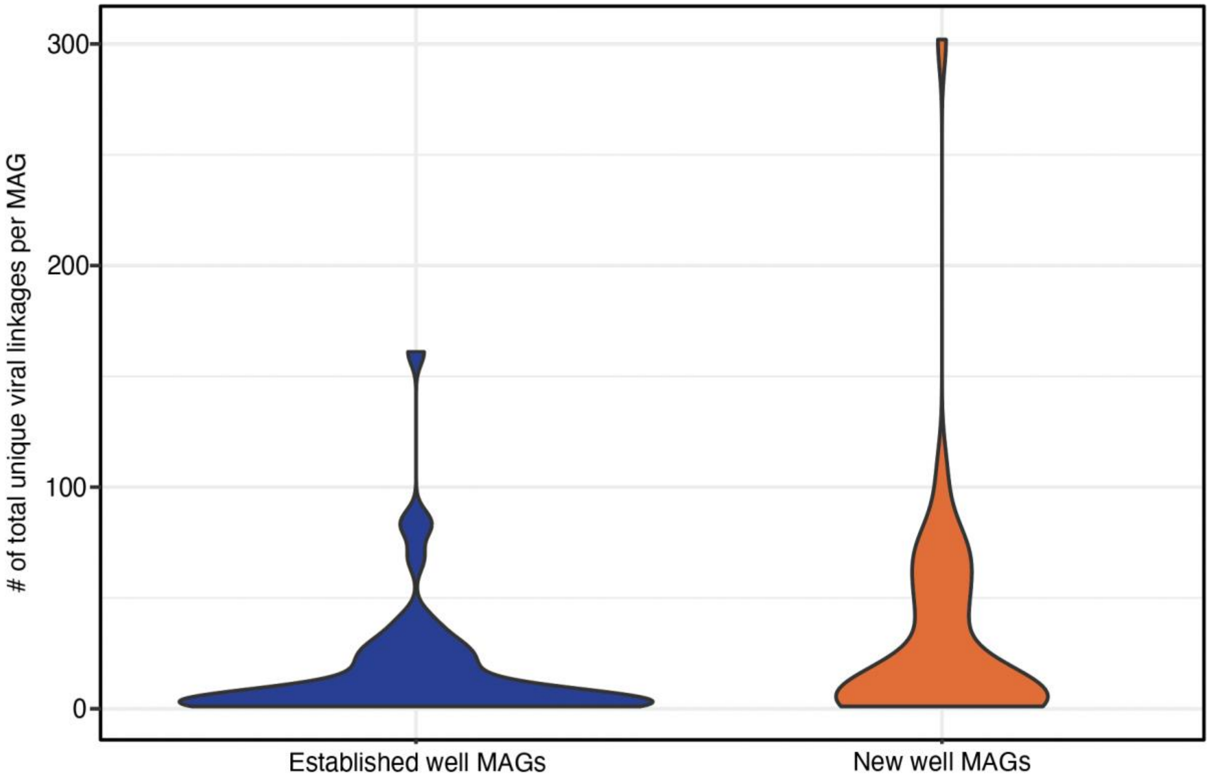


**Figure S13.** Distribution of total number of linkages, per MAG, to unique vMAGs. Linkages were made using paired CRASS recovered spacers as well as Geneious identified spacers and therefore viral linkages may have been made via spacers recovered at many different timepoints, but distribution shows the total per MAG.

### Supplementary Discussion

#### ***Evidence that viral predation is likely occurring in the DJ Basin***

We observed higher host (bacterial and archaeal) alpha diversity in the established wells relative to the new wells (*Wilcox, p=5.563e-06*) (Figure 1). This trend contrasts findings from previous fractured shale studies that reported a rapid decrease in microbial diversity following the hydraulic fracturing process (1,2). More broadly, host alpha diversity in the DJ Basin was also higher than many other fractured shale ecosystems studied to date (3–6) and similar to those reported previously for produced fluids from the DJ Basin (7). This may be due to the lower salinity in the DJ Basin exerting less selective force on colonizing microorganisms and thus allowing for more taxonomically diverse communities (8–10). Trends in viral alpha diversity closely mirrored those observed in the host communities, with higher viral diversity in the established wells. Given the reliance of viruses on their hosts for replication, alpha diversity trends of host and viral communities were strongly and positively correlated through time, providing evidence for ongoing viral predation and proliferation in all six wells (*Spearman Rho: old=0.71, 0.96, 0.076 new=0.6, 0.66, 0.91*).

We quantified temporal changes in community structure using Bray-Curtis dissimilarity values to investigate how these taxonomically diverse communities fluctuated through time (Figure S4). In this analysis, higher dissimilarity values indicate greater change in community composition relative to the previous timepoint. In all wells, shifts in host and viral communities were strongly and positively correlated, often mirroring one another in their dynamics (Figure S4). Viruses depend on their hosts for replication, yet host populations are often impacted as a direct result of this proliferation. Thus, the strong relationship observed here suggests that host and viral communities are continually changing, and viral predation is likely occurring. Interestingly, we observed a stronger relationship between host and viral communities in the established wells compared to the new wells. Strong correlations between host and virus have been previously shown in less diverse shale microbial communities, specifically with *Halanaerobium* and associated viruses in the Appalachian Basin (11). Here we report similar trends in a more taxonomically diverse host community over an extended timeseries and in six distinct shale wells.

***Relationships between number of spacers with time and host coverage are obscured at higher taxonomic levels.***
In general, the number of spacers associated with a given MAG had a strong positive relationship with MAG coverage, a proxy for relative abundance in the community. This connection provides further evidence that encoding a CRISPR system and accumulating spacers as a result of viral interactions may support the host's overall success in the ecosystem. This trend was obscured at higher taxonomic levels, however, highlighting the importance of genome-resolved analyses (Figure S5). It is worth noting that not all MAGs shared this same relationship with number of spacers and coverage; in the deep terrestrial subsurface other physicochemical parameters (e.g., temperature, salinity) likely play a key role in environmental filtering and shaping the persisting microbial community.

Although these results highlight the importance of CRISPR-mediated viral defense in this subsurface ecosystem, the number of spacers associated with many MAGs varied greatly through time, with few MAGs exhibiting a steady increase in total number of spacers. In contrast to the relationship with MAG coverage, correlations between number of spacers and time varied most at the genome-level yet was more constrained to a weakly positive average relationship at the phylum-level (Figure S5). The greater variability measured at the genome level is due to some host populations exhibiting a strong positive relationship between number of spacers and time while many others do not. This could be due to myriad reasons, such as the overall temporal dynamics of the host population, stochastic loss and gain of spacers through time, bioinformatic limitations, or perhaps preferential use of other viral defense systems besides CRISPR.

***Viral defense systems, besides CRISPR, encoded in MAGs***

Most DJ Basin MAGs also contained other defense systems besides CISPR-Cas, and these systems were identified using DefenseFinder (12). There was a greater diversity of different viral defense systems encoded in the established wells compared to the new wells, with 41 and 34 different systems detected across all MAGs, respectively. In the established wells, we observed viral defense systems other than CRISPR across 84 of 105 MAGs (80%), averaging 4.2 viral defense systems per MAG. In the new wells, we observed viral defense systems across 75 of 97 MAGs (77%) averaging 3.6 systems per MAG. In both established and new wells, the most common defense system observed was a restriction modification defense mechanism. Most of the MAGs that contained a CRISPR array also tended to encode another defense system in both the new and established wells, with 81% and 78% of MAGs respectively. Additionally, many of the MAGs without a detected CRISPR array also encoded a different viral defense system, with 32 of 44 (72%) of MAGs without CRISPR in the new wells encoding a different viral defense system, and 29 out of 35 MAGs (82%) in the established wells. In the established wells, this left only 6 MAGs (of 105 total) that we did not detect any viral defense mechanism, though this could be due to partial assembly and/or completeness of the genomes. In the new wells, there were 12 of 97 MAGs without a detectable CRISPR or other viral defense system. Together, this provides further evidence that viral predation is likely occurring in this fractured shale ecosystem, and that there is likely a benefit to the host for encoding viral defense mechanism(s).

***Viral lifestyle likely not driving virus-host co-occurrence trends***

Viruses have many different ‘lifestyle’ types that can have different effects on an individual host cell, host population, and overall microbial community. For the purpose of our study, we simplified these different lifestyle types into only lytic and temperate to determine if a temperate or prophage lifestyle may be driving the co-existence trends we observe. If this was the case, more instances of a successful interaction or ‘co-occurrence’ would be found if the virus was integrated into the host. For this analysis we considered all MAGs linked with all individual vMAGs, which resulted in 815 different vMAGs linked across 48 MAGs in the established wells, and 1295 vMAGs linked across 42 different MAGs in the new wells. We surveyed our viral genomes for integrase genes – which may indicate a temperate lifestyle – and found that a small proportion of our viral community may have the potential for that lifestyle. Of our total 2,176 viral genomes, we observed 192 vMAGs contained an integrase gene (annotated via KEGG), although fewer of these viral genomes were recovered from the new wells (*n=65*) compared to the established wells (*n=126*). Additionally, 11 of the 192 potentially temperate viruses were not present (via our requirements for metagenomic read recruitment) in any of the 78 timeseries samples. Only a very small proportion of vMAGs (<5 in each well) exhibited temporal patterns in their coverages that nearly matched their hosts’ coverage, indicating that though they may have the potential for a temperate lifestyle, their paired dynamics are not solely responsible for the trends we observe in host-viral co-occurrences.

### Supplementary Methods

***Additional details on hydraulic fracturing, subsurface shale ecosystems, and specifics of sampling fractured shales.***

We sampled the produced and formation fluids from six hydraulically fractured shale wells from the Niobrara formation within the Denver-Julesburg (DJ) Basin located in eastern Colorado. The Niobrara shale formation consists of three benches that are located approximately 1890-1950 meters deep in the subsurface with a downhole temperature measuring approximately 112°C. Hydraulically fractured shale ecosystems are anthropogenically formed for the recovery of oil & gas that occur throughout North America and elsewhere. Hydraulic fracturing, or "fracking" occurs by first drilling vertically to the target shale formation and then laterally through it to enhance oil & gas recovery. Next, water, sand, and chemicals are injected at high pressure to create fractures & fissures thereby releasing the otherwise trapped oil & gas, which flows back to the earth's surface with water. Over the development of a well, this water transitions from mostly injected water to mostly formation water. Inputs used are not sterile and therefore introduce many microbes into this newly formed ecosystem during the hydraulic fracturing process. A subset of these microorganisms are able to survive the deep subsurface shale physiochemical conditions (elevated salinity, temperature, and pressure) and colonize and persist in this ecosystem throughout the lifetime of the well. Deep subsurface shale formations are likely extremely low biomass prior to hydraulic fracturing, as shales are compact rock formations that are often characterized by limited water, limited pore connectivity within the formation, and very small pore sizes that are often smaller than the average bacterial cell.

In this study we sampled early fluids directly from the well head, however most fluid samples (those named with 'DJKA#') were collected from the corresponding separator tanks that are unique to each well. Separator tanks separate the oil, gas, and water portions of the flowback fluid. Therefore, MAGs were recovered from produced fluids collected from the separator tank or well head for each well, though for brevity we refer to these MAGs as simply recovered from the well. The target bench for each well was either Niobrara bench B (DJB-1, DJB-3, and DJB-4) or Niobrara bench C (DJB-2, DJB-5, and DJB-6) with all wells having a lateral length of either one or two miles. The new wells began 'producing' in October 2018 while the established wells had been in production since January of 2016.
